# Predicting the effect of non-coding mutations on single-cell DNA methylation using deep learning

**DOI:** 10.1101/2024.09.03.611114

**Authors:** Zhe Liu, An Gu, Yihang Bao, Guan Ning Lin

**Author notes:** Authors to whom correspondence should be addressed. (Guan Ning Lin).

## Abstract

Predicting the effects of non-coding mutations on DNA methylation is crucial for advancing our understanding of gene expression, epigenetic inheritance, and its role in disease mechanisms. Current methods lack the capability to predict the impact of non-coding mutations on DNA methylation at single-cell resolution and long range, while remain challenges in tracking SNP influences throughout disease progression. Here, we introduce Methven, a deep learning-based framework designed to predict the effects of non-coding mutations on DNA methylation at single-cell resolution, to overcome the challenges. Methven integrates DNA sequences and ATAC-seq data, employing a divide-and-conquer approach to handle varying scales of SNP-CpG interactions. By leveraging a pretrained DNA language model, Methven accurately predicts both the direction and magnitude of methylation changes across a 100kbp range with a lightweight architecture. The evaluation results demonstrate the superior performance of Methven in prioritizing functional non-coding mutation, model interpretability, and its potential for revealing personalized mutation-disease associations.

## Introduction

The regulation of gene expression through DNA methylation is central to understanding cellular differentiation and disease pathogenesis[1, 2]. Despite the crucial role of DNA methylation, current strategies to analyze genetic variants, particularly those identified by genome-wide association studies (GWAS), remain limited[3]. GWAS efficiently identify numerous variants associated with diseases and traits; however, they often do not resolve the functional consequences of these variants[4, 5]. Traditionally, the analysis of genetic variants has concentrated on direct genomic signals, such as transcription factor binding and histone modifications[6-8]. These methods, while informative, frequently overlook the subtle yet critical effects of DNA methylation. This oversight can obscure deeper insights into cellular responses and developmental processes that are vital for a comprehensive understanding of gene regulation.

CpGenie[3], a tool specifically designed to predict the impact of non-coding variants on DNA methylation, illustrates some of these challenges. While it advances the field by focusing on the methylation impacts of sequence variants, its prediction reach is constrained by a limited receptive field of 500 base pairs (bp). This narrow scope restricts its utility in conditions where the regulatory effects of a mutation span broader genomic regions.

In contrast, tools like Enformer[9] and DeepSea[10], which annotate DNA sequences for potential functional impacts, primarily offer static predictions with longer receptive field. These models, though valuable, do not adequately capture the temporal dynamics of epigenetic modifications that occur in response to environmental changes or disease progression. They also tend to focus on broader aspects of gene regulation without providing the resolution needed to dissect the specific effects of individual non-coding variants on methylation patterns.

Furthermore, there is a notable gap in the availability of tools that can predict the effects of non-coding DNA mutations on methylation at a single-cell resolution. Such capabilities are essential for personalized medicine, as they would allow for the detailed characterization of epigenetic variability at the level of individual cells.

To address these shortcomings, we developed Methven, a deep learning framework designed to predict the effects of non-coding mutations on DNA methylation at single-cell resolution. Methven integrates DNA sequences with ATAC-seq data using a divide-and-conquer strategy that addresses SNP-CpG interactions across variable distances up to 100kbp with a lightweight architecture. The framework supports dual tasks: classification to determine the direction of methylation change and regression to quantify its magnitude, enhancing predictive accuracy. We show that Methven outperforms existing methods in cell-specific predictions, while offering a dynamic solution that adapts to the cell-specific and progressive nature of epigenetic changes, which is crucial for understanding disease mechanisms and advancing personalized medicine.

The efficacy of Methven is demonstrated through various validations. Ablation studies confirm the necessary of its components, enhancing the model’s accuracy and robustness. Visualization of representation capabilities via t-SNE illustrates Methven’s nuanced understanding of genomic interactions. Further, its performance on external cell types like monocytes underscores its generalizability and potential for personalized medicine. Lastly, hidden state analyses highlight Methven’s ability to discern and utilize complex regulatory patterns effectively.

## Results

### Overview of Methven

Methven is designed to predict the impact of non-coding mutations on all methylation sites within a 100kbp range upstream and downstream at single-cell resolution, by training on methylation quantitative trait loci (meQTL) data[11] that both DNA sequences and ATAC-seq are available. Training on meQTL data minimizes the risk of model bias caused by false-negative SNPs (SNPs that affect CpG sites but are not statistically captured), while the inclusion of an ATAC-seq input branch ensures that the model retains the potential for generalization and fine-tuning across other cell types or tissues.

The architecture of Methven consists of two main components (Fig. 1): (1) preprocessing of labeled SNP-CpG pairs to generate embeddings suitable for training, and (2) a deep learning module designed to perform both classification and regression tasks.

**Fig 1.**
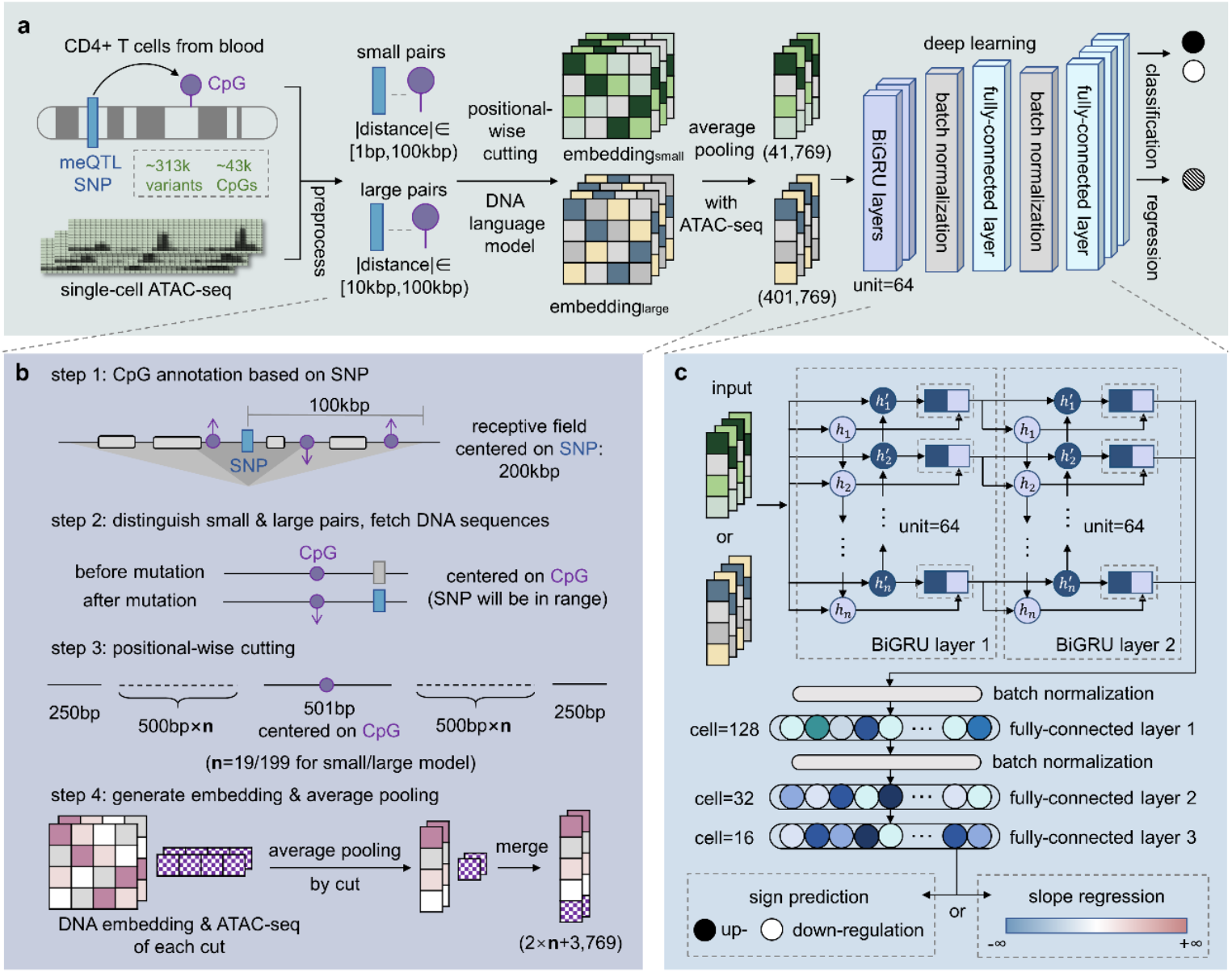
Overview of Methven. (a)Prediction of mutation impacts on methylation using DNA sequences and single-cell ATAC-seq data. In the data collection phase, meQTL data from CD4+ T cells were used to label the impact of non-coding mutations on CpG sites. A divide-and-conquer strategy was employed, segmenting the dataset into small pairs and large pairs based on the distance between SNPs and CpG sites. DNA sequences after positional-wise cutting were fed into a DNA language model. The generated DNA embeddings and the corresponding ATAC-seq data were averaged and concatenated within each cut unit. The concatenated embeddings were finally input into the deep learning model designed to perform both classification and regression tasks. (b) Illustration of the preprocessing pipeline. For each SNP, CpG sites within 100kbp upstream and downstream were identified based on methylation changes. SNP-CpG pairs within 10kbp formed the small dataset, while those between 10kbp and 100kbp formed the large dataset. DNA sequences and corresponding ATAC-seq data were extracted with the CpG site centered. The sequences, both pre- and post-mutation, were then positionally cut around the CpG site, input into a DNA language model to obtain embeddings. Finally, the DNA embeddings from each cut and the ATAC-seq data were average pooled and concatenated. (c) Details of the deep learning module. The concatenated embeddings are fed into two stacked Bidirectional Gated Recurrent Unit (BiGRU) modules, followed by batch normalization layers and fully connected layers. The classification and regression tasks are handled separately: the classification task predicts the direction of the SNP’s impact on CpG methylation levels (upregulation/downregulation), while the regression task estimates the magnitude of this impact (slope).

First, during data preprocessing, we annotated CpG sites within a 100kbp range upstream and downstream of each SNP that were affected by the mutation and constructed a comprehensive dataset (Fig. 1b). The meQTL data for CD4+ T cells were obtained from the meQTL EPIC Database[12], while the corresponding ATAC-seq data were sourced from the EpiMap Repository [13].

To address the varying nature of SNP-CpG interactions, we employed a divide-and-conquer strategy. Recognizing that SNPs in close proximity to CpG sites may exhibit more direct and stronger effects, while those at greater distances might involve more complex long-range regulatory mechanisms, we divided the dataset into small pairs (distance < 10kbp) and large pairs (distance between 10kbp and 100kbp). Independent models were trained for each subset, allowing each model to focus on capturing the specific features relevant to their respective scales. For small pairs, DNA sequences and ATAC-seq data within a 10kbp range centered on the CpG site were utilized as initial features, excluding SNPs with insufficient sequence length. For large pairs, this range was extended to 100kbp.

Next, to enhance the representation of DNA sequences, we selected DNABert2[14] as the language model for generating pre-trained DNA embeddings. Due to the input sequence length limitations of DNABert2, we developed a positional-wise cutting strategy. This approach ensures that each segment remains within the model’s input limits while centering the CpG site as the critical focal point, thereby maximizing the retention of key information. For both pre- and post-mutation sequences, the DNA embeddings and ATAC-seq data for each segment were average pooled and concatenated.

Finally, the concatenated embeddings were input into Methven’s deep learning module. This module comprises two stacked Bidirectional Gated Recurrent Unit (BiGRU) layers[15], followed by batch normalization layers[16] and fully connected layers[17]. In this study, we designed two independent tasks: a classification task to predict the direction of the SNP’s impact on CpG methylation levels (up-regulation/down-regulation), and a regression task to estimate the magnitude of this impact (slope). By separating these tasks, Methven enables the model to focus on each predictive goal independently, minimizing task interference and achieving higher predictive accuracy, particularly in determining the direction of the methylation impact.

### Evaluating performance on intra-dataset

We first validated Methven’s prediction performance on the internal test set (Fig. 2a-b). In the classification task, predicting the direction of the SNP’s impact on CpG methylation without considering the magnitude of the effect amplifies the differences between the two, thereby reducing potential interference from samples with smaller absolute slope values that might obscure the directional judgment. On a test set comprising 1,988 small SNP-CpG pairs, the Methven classification model achieved an ACC of 0.920 and an AUC of 0.969. For a test set of 3,032 large SNP-CpG pairs, the ACC was 0.837, and the AUC was 0.918. These results demonstrate the robust predictive capability of the Methven classification model, with a receptive field extending up to 200kbp.

**Fig 2.**
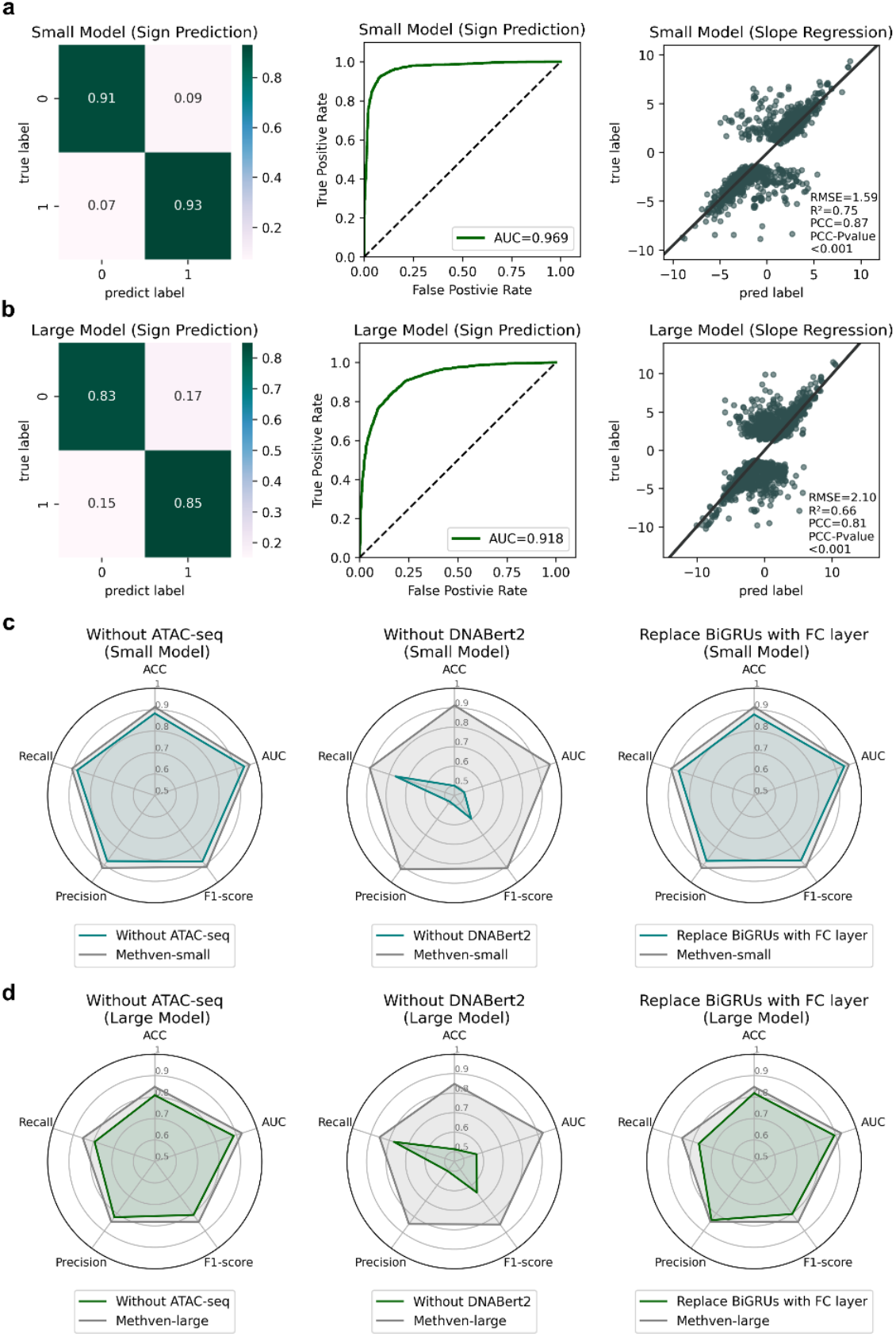
Benchmarking and robustness evaluation of Methven on intra-dataset. (a) Performance of Methven on the small dataset. The classification task is evaluated using a confusion matrix, ROC curve, AUC, accuracy (ACC), recall, precision, and F1-score. The regression task performance is quantified by RMSE, R^2^, and PCC, with the significance of PCC assessed under the condition of P < 0.001. (b) Performance of Methven on the large dataset. Evaluation metrics are applied similarly to those in (a). (c) Ablation study on the small dataset. Four ablation experiments were conducted: removing the input ATAC-seq data, removing the DNA embedding, and replacing the BiGRU layers with fully connected layers. Grey lines represent the performance of the full Methven model, while green lines depict the performance of the ablated models. (d) Similar to (c), with the same ablation experiments conducted on the large dataset.

However, predicting only the direction of the SNP’s impact on CpG methylation is insufficient. Therefore, we trained additional independent models to regress the magnitude of this impact, specifically the meQTL slope. On the test set, the Methven-small model achieved an RMSE of 1.59 and a PCC of 0.87 (t-test p < 0.001), while the Methven-large model recorded an RMSE of 2.10 and a PCC of 0.81 (t-test p < 0.001). Although the regression results displayed two clusters along the diagonal, likely due to the unique characteristics of the training data, the statistical evaluation metrics indicate that Methven can capture the extent of the SNP’s impact on CpG methylation. This suggests that when the predicted absolute slope value is small, the SNP-CpG pair likely has minimal association, making further assessment of the impact direction unnecessary.

The ablation experiments demonstrated that each critical component of Methven independently contributes to its overall performance. These key components include the ATAC-seq input, DNA embeddings, and the BiGRU layers within the model architecture. We conducted a series of experiments where we removed the ATAC-seq input, excluded the DNA embeddings, and replaced the BiGRU layers with fully connected layers. As shown in Fig. 2c-d, the performance decline following the removal of these essential components underscores the validity of Methven’s design.

To further verify the effectiveness of the encoder of Methven, we utilized t-SNE algorithm[18] to visualize the sample distribution based on the initial feature (the both pre- and post-mutation DNA embedding) and representation generated via Methven, respectively. All embeddings and representations were mapped into a two-dimensional space. To visualize the sample distribution, SNP-CpG pairs were colored according to the SNP’s effect on methylation (Fig. 3). We observed that in both the classification and regression tasks, SNP-CpG pairs with different labels or slopes were completely intermixed in the two-dimensional space of the initial characterization. However, in the Methven representation space, SNP-CpG pairs were separated according to their classification labels (Fig. 3a) and were distributed in an orderly manner according to slope values in the regression task (Fig. 3b). These results indicate that Methven is able to efficiently generate high-quality representation vectors for mutation effect prediction and maintain a consistent performance in different tasks.

**Fig 3.**
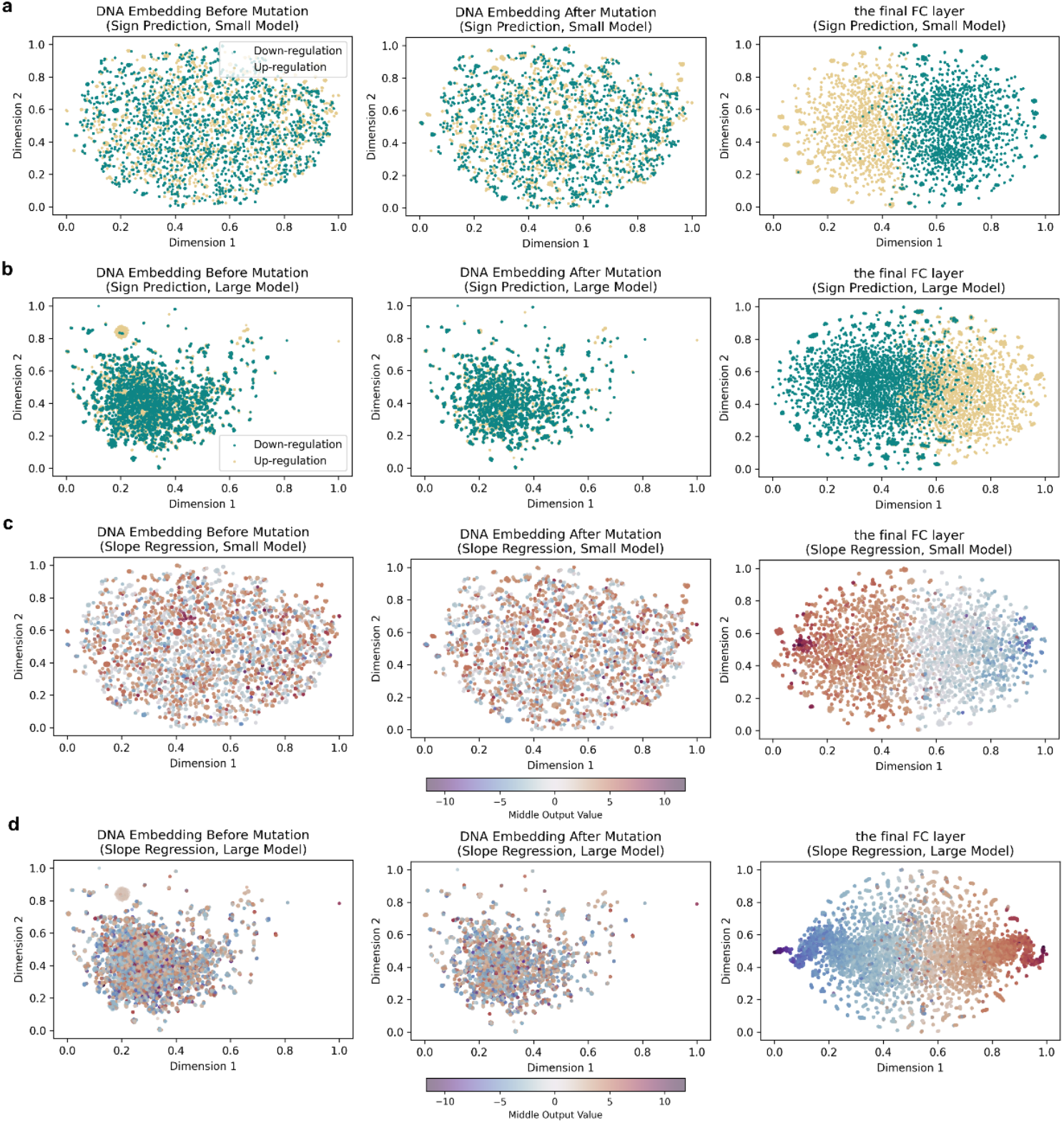
Visualization of representation ability of Methven. (a) Visualization of representational abilities in the Methven small model for the classification task. t-SNE was used to perform dimensionality reduction and visualization on the DNA embeddings both pre- and post-mutation, as well as on the outputs from the Methven model after the deep learning module. Green points represent SNP-CpG pairs where the CpG methylation level increases, while yellow points represent pairs where the CpG methylation level decreases. (b) Similar to (a), with t-SNE applied to the large dataset model. (c) Visualization of representational abilities in the Methven small model for the regression task. t-SNE was used to perform dimensionality reduction and visualization on the DNA embeddings both pre- and post-mutation, as well as on the outputs from the Methven model after the deep learning module. Points are colored with a gradient from blue to red, representing slope values from low to high. (d) Similar to (c), with t-SNE applied to the large dataset model.

### Comparison with external methods demonstrating Methven’s superiority in non-coding mutation effect prediction

Methven can predict the impact of SNPs on all CpG sites within a 100kbp range upstream and downstream, whereas the previous state-of-the-art model, CpGenie[3], was limited to a 500bp range. To comprehensively demonstrate Methven’s effectiveness and robustness in predicting the effects of non-coding mutations on methylation, we compared it with existing external tools. CpGenie is currently the best-performing and the only method specifically designed for predicting the impact of mutations on methylation. To ensure a fair comparison, we extended the CpGenie architecture to accept data with SNP-CpG distances up to 100kbp.

Enformer[19] was originally designed to predict the impact of non-coding mutations on gene expression but also excels at generating functional annotations of DNA sequences. On the other hand, Methven’s features are derived from large-scale pretraining on DNA sequences to learn semantic representations. Comparing Methven with Enformer allows us to evaluate which annotation approach—functional or semantic—provides stronger predictive capabilities for the task of assessing the impact of non-coding mutations on methylation.

The comparison was conducted using ten-fold cross-validation on the classification task. To fairly assess the representational power of different methods, we extracted the embeddings from the penultimate layer of each model (the embeddings used for final classification, representing the highest-level features learned by the model). We then trained a decision tree with default parameters on these embeddings and evaluated the models on an internal test set (Fig. 4a, Fig. S3).

**Fig 4.**
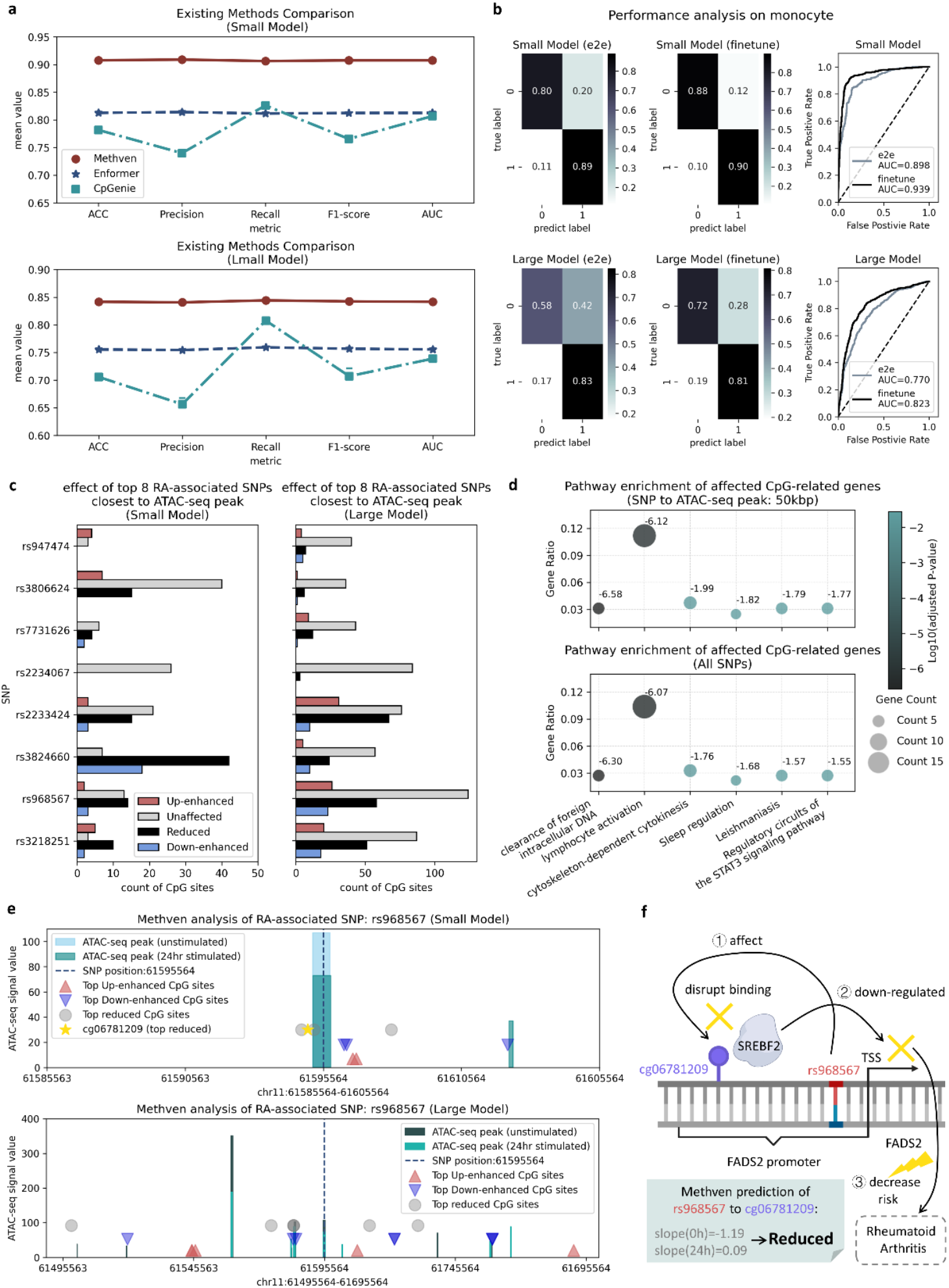
External validation of Methven on existing methods, new cell type, and disease-associated SNPs. (a) Comparison of ten-fold cross-validation performance between Methven, Enformer, and CpGenie on the classification task. The metrics used for comparison include ACC, Precision, Recall, F1-score, and AUC. To ensure a fair comparison of the models’ ability to learn the relationship between SNPs and CpG sites, embeddings from the layer preceding the output layer of each model were extracted, and a decision tree with identical parameters was trained on these embeddings. (b) Methven’s performance on monocyte single-cell meQTL datasets. The experiments involved two approaches: end-to-end (e2e) training of Methven directly on the monocyte dataset, and fine-tuning Methven pre-trained on the CD4+ T cell dataset. (c) Analysis of rheumatoid arthritis (RA)-associated SNPs using the Methven small regression model. The SNPs predicted were selected from GWAS analysis as RA-associated SNPs. Red bars indicate SNPs where the absolute difference in slope between case and control SNPs is greater than 0.5, with the control SNP having a positive slope, suggesting an up-enhancement of methylation impact in RA cases. Blue bars represent SNPs where the absolute difference in slope is greater than 0.5, with the control SNP having a negative slope, indicating a down-enhancement of methylation impact in RA cases. Grey bars indicate SNPs where the absolute difference in slope is less than 0.5, suggesting little association with RA in terms of methylation impact. Black bars denote SNPs where the absolute slope in RA cases is smaller than in controls, indicating a reduced impact on methylation in RA conditions. (d) Pathway enrichment results for genes corresponding to Methven-predicted affected CpGs. To compare the impact of varying signal-to-noise ratios, the SNPs within a 50kbp range of the ATAC-seq peak were analyzed separately and compared to the results from all SNPs. (e) Visualization of RA-associated SNP analysis for the Methven. rs968567 has been reported as a key SNP for RA. For the different categories of SNPs identified in (c), top-ranking CpG sites were selected as Top CpG sites, showing that most are located near ATAC-seq peaks. (f) Case study of Methven’s prediction for the rs968567 to cg06781209 interaction. cg06781209 is a binding site for the transcription factor SREBF2. rs968567 in the promoter region of the FADS2 gene alters DNA methylation, disrupting the binding of SREBF2 and downregulating FADS2 expression, thereby reducing RA risk. In Methven’s predictions, the absolute value of the unstimulated slope is greater than that of the case slope, indicating that in RA, the CpG site is less influenced by the SNP. This suggests a reduced ability of the SNP to modulate CpG methylation, thereby failing to suppress RA as effectively.

We found that Methven outperformed all other methods on both the small and large datasets across all evaluation metrics, with consistently balanced recognition of positive and negative samples (Methven small model: mean ACC=0.908, mean AUC=0.908; Methven large model: mean ACC=0.842, mean AUC=0.842, Table S5). In contrast, while Enformer also achieved balanced recognition of positive and negative samples, its overall performance was inferior to Methven, likely because the functional annotation features learned during pretraining of Enformer are not fully suited for methylation tasks. CpGenie, which uses OneHot encoding for DNA sequences and convolutional neural networks (CNN) for representation learning, struggled with longer sequence lengths (> 500bp reported), resulting in less stable and poorer performance. This result demonstrates the effectiveness and robustness of Methven in predicting the impact of non-coding mutations on DNA methylation.

### Validation across external cell type

The effect of mutations can vary across different cellular environments. To explore Methven’s potential for generalization to other cell types, we applied the Methven classification model to a dataset comprising monocyte single-cell meQTLs. This dataset was downloaded from the EPIGEN MeQTL Database[12], with corresponding ATAC-seq data sourced from the EpiMap Repository [13]. After preprocessing, the entries used for training and testing were approximately one-third of those in Methven’s internal dataset (Table S1, Table S8).

We first trained Methven directly on the monocyte external validation dataset using an end-to-end (e2e) approach. We observed solid classification performance on both the small pairs and large pairs (AUCs of 0.898 and 0.770, respectively, Fig. 4b, Table S6). This indicates that Methven has a certain level of potential to generalize to other cell types or tissue types, provided that corresponding ATAC-seq data is available.

Next, we attempted fine-tuning Methven based on the pre-trained model from CD4+ T cells. We found that the fine-tuned model showed a slight improvement in performance over the end-to-end training (AUCs of 0.939 and 0.823, respectively, Fig. 4b), which could be attributed to the larger number of entries in Methven’s internal dataset, allowing it to learn more intrinsic relationships between meQTLs and ATAC-seq. This suggests that Methven has the potential to serve as a generalized pre-trained model, particularly as the amount and the cell/tissue type of training data increases over time.

### Exploring potential of uncovering mutation-disease mechanisms through analyzing disease-associated SNPs

In real-world circumstances, studies on the effects of mutations often involve exploring disease mechanisms[20, 21]. Thanks to the incorporation of ATAC-seq inputs to capture epigenetic states, Methven enables the analysis of mutations occurring in the same cell type under different disease processes.

To explore Methven’s potential in elucidating the connection between mutations and disease mechanisms, we selected SNPs highly associated with rheumatoid arthritis (RA) through GWAS analysis[22]. For training, we used ATAC-seq data from CD4+ T cell lines, with cells stimulated for 24 hours with anti-CD3/CD28 serving as the case ATAC-seq, and unstimulated cells as the control ATAC-seq[23]. To enhance the signal-to-noise ratio, we filtered SNPs located within 1kbp upstream and downstream of the control ATAC-seq peak regions, resulting in 8 remaining SNPs, and annotated all CpG sites within a 100kbp range upstream and downstream of these SNPs.

Additionally, we mapped the predicted affected CpGs to the nearest TSS within the genome. To enhance the signal-to-noise ratio, we conducted pathway enrichment analysis for the affected CpGs-related genes observed by all SNPs and the SNPs located within a 50kbp range of the ATAC-seq peaks. We then identified the pathways with significant adjusted p-values that were common to both groups, finding that ‘clearance of foreign intracellular DNA’ and ‘lymphocyte activation’ were both highly significant pathways (Fig. 4d). The clearance of foreign intracellular DNA is reported closely linked to the pathogenesis of RA through its role in the immune response and inflammation[24], while the signaling lymphocytic activation molecule family (SLAMF) may influence RA pathogenesis by participating in inflammation mediated by infiltrating immune cells[25]. The identification of RA-associated CpGs influenced by SNPs and recognized by Methven, which align with pathways known to be involved in RA pathogenesis, demonstrates Methven’s capability to enhance the understanding of the role of ‘mutations affect methylation levels’ in disease mechanisms.

We then used the Methven regression model to predict the impact of these SNPs and their corresponding ATAC-seq data in both case and control samples. Based on the differences in slopes between the two conditions, we categorized the annotated CpG sites into four groups: those with mutation impact of methylation enhanced by disease occurrence (Up-enhanced), those with mutation impact of methylation negatively enhanced (Down-enhanced), those with mutation impact of methylation reduced impact (Reduced), and those with mutation impact of methylation unaffected by the disease (Unaffected). We found that Methven could distinguish among these four categories of CpG sites, addressing a gap left by other methods in this functional area (Fig. 4c).

To further illustrate Methven’s capabilities, we used rs968567—a SNP proven to be highly associated with RA—as an example, showcasing the relationship between the SNP, case and control ATAC-seq peaks, and the representative positions of the four categories of CpG sites relative to the SNP (Fig. 4e, Table S7). We observed that the Top CpG sites were mainly clustered around the ATAC-seq peak, regions more likely to contain functional regulatory elements such as enhancers and promoters. This clustering may explain why these CpG sites are more profoundly impacted by the SNP and exhibit greater changes due to disease occurrence.

Notably, cg06781209 is a binding site for the transcription factor SREBF2, and rs968567, located in the promoter region of the FADS2 gene, has been reported to alter the methylation level of cg06781209[26]. This alteration disrupts the binding of SREBF2, downregulating FADS2 gene expression and subsequently reducing RA risk[26]. In other words, changes in the methylation level of cg06781209 associated with the suppression of the effects of rs968567, thereby influencing RA risk. Methven’s predictions revealed that the CpG methylation levels affected by the 24h stimulated SNP showed less fluctuation compared to the unstimulated SNP (Fig. 4f), suggesting that the SNP’s ability to influence transcription factors—and consequently RA—was diminished. The prediction results indicat that in RA, the CpG site is less influenced by the SNP, aligning with the aforementioned findings.

These results demonstrate that Methven can assist in determining whether the pathogenicity of disease-associated SNPs is driven by “mutations affecting methylation levels,” thus providing insights into the impact of individual mutations on disease risk and progression, and supporting the development of personalized treatment strategies.

### Investigating model interpretability through hidden state analysis

Non-coding DNA typically contains numerous functional regulatory regions, and SNPs located near these regions are more likely to impact gene expression and methylation. To explore Methven’s ability to understand and utilize sequence features, we aligned the hidden states of the BiGRU layers with the DNA sequences and analyzed them based on different functional regions, as well as whether the DNA bases were located within these functional regions. We obtained annotations for eight types of functional regions from the UCSC Genome Browser: active promoter, strong enhancer, transcriptional transition, transcriptional elongation, insulator, heterochrome, repressed region, and repetitive element/copy number variation.

We observed that the activation values of the BiGRU layer’s hidden states differed significantly between functional regulatory regions and non-functional regions (Mann-Whitney U test, P<0.005, Fig. 5). This finding suggests that the model likely recognizes and leverages the information from these critical regions, contributing to its superior performance in the classification task. This indicates that the model not only excels in overall performance but also demonstrates an advanced capability in understanding and utilizing sequence features.

**Fig 5.**
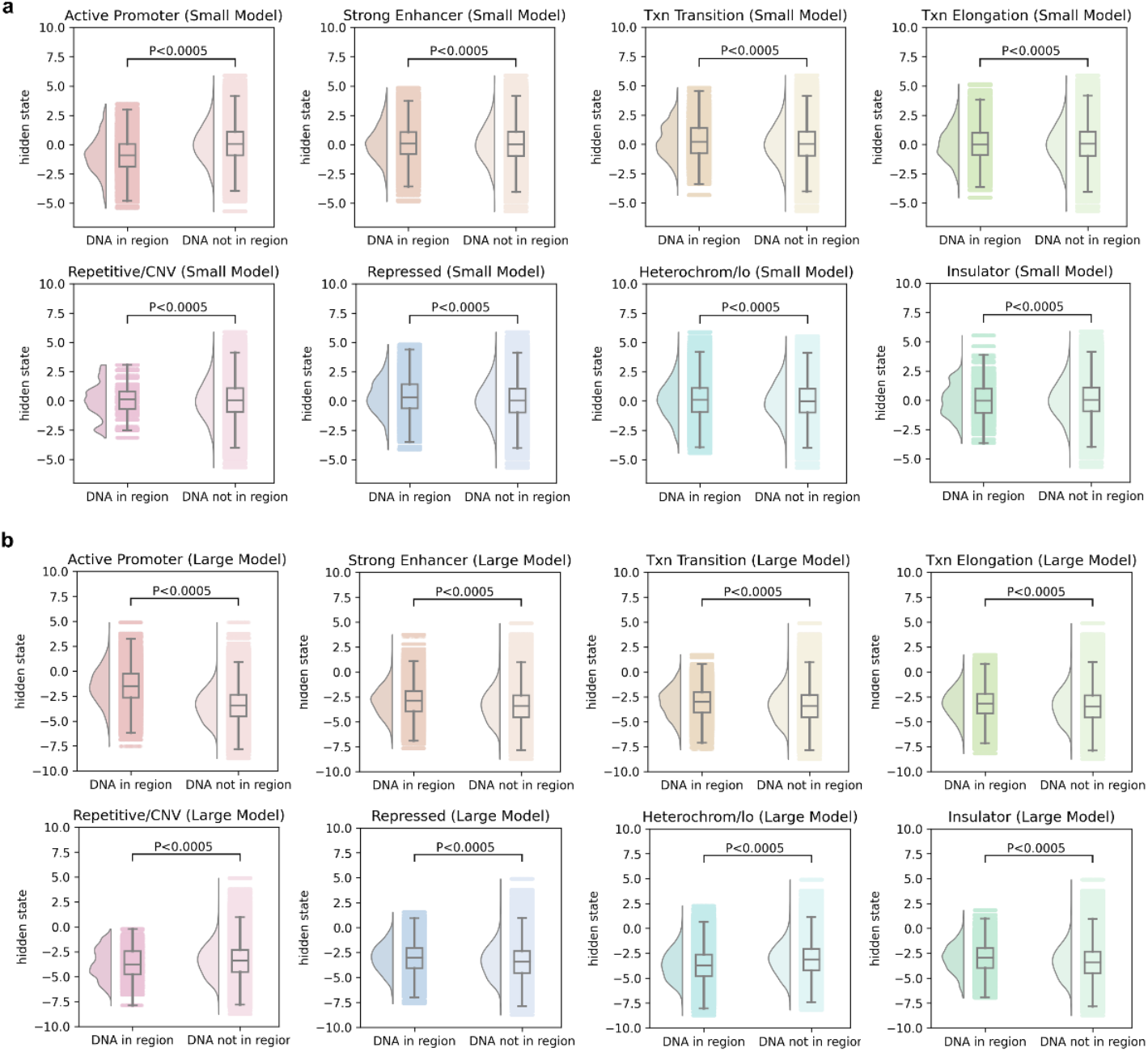
Hidden state analysis of Methven. (a) Differences in the hidden states of stacked BiGRU layers in the Methven small model between functional and non-functional regions. Eight functional regions were annotated using the UCSC Genome Browser and mapped onto the DNA sequences of the input SNP-CpG pairs (Txn Transition: transcriptional transition, Txn Elongation: transcriptional elongation, Repetitive/CNV: repetitive element/copy number variation, Repressed: repressed region, Heterochrom/lo: heterochrome). The differences between functional and non-functional regions were assessed using the Mann-Whitney U test. (b) Similar to (a), with the analysis conducted using the Methven large model.

Moreover, we found distribution differences in the hidden states between the Methven small and large models, suggesting that the model learns different regulatory patterns for short-range and long-range interactions. This observation underscores the potential differences in regulatory mechanisms at varying distances and further validates the rationale behind Methven’s strategy of training separate models for different sequence lengths.

## Discussion

Despite notable advances in GWAS in identifying genetic variants linked to diseases and traits, pinpointing causal variants and deciphering their pathogenic mechanisms remain formidable challenges. The state-of-the-art method of predicting the effect of non-coding mutations on DNA methylation, CpGenie, while pioneering in predicting non-coding variant effects on DNA methylation, is confined by their limited receptive fields and lack dynamic response capabilities to changing cellular environments throughout disease progression. Similarly, while tools like Enformer and DeepSea provide function annotation and static predictions from DNA sequences, they fall short in adapting these predictions to the dynamic, cell-specific contexts critical during disease states.

Methven addresses these gaps by integrating DNA sequences with ATAC-seq data, employing a divide-and-conquer strategy to handle SNP-CpG interactions across extensive genomic ranges, up to 100 kbp. This integration, supported by the utilization of the advanced DNA language model DNABert2, allows Methven to predict both the direction and magnitude of methylation changes at a single-cell resolution with a lightweight model structure. Methven’s architecture, which supports dual tasks of classification and regression, minimizes task interference and enhances the model’s robustness.

The validation of Methven on diverse cell types, including monocytes, demonstrates its generalizability and potential for personalized medicine. However, Methven is not without limitations. Its performance, while superior in the conditions tested, may vary with different epigenomic landscapes not represented in the training data. Furthermore, the computational demands of processing extensive genomic ranges and the need for high-quality, cell-specific ATAC-seq data may limit its application in less resourced settings.

Future improvements for Methven could include expanding its training datasets to encompass a wider array of cell types and conditions to enhance its adaptability and accuracy. Additionally, integrating other forms of epigenetic data, such as histone modification patterns, could provide a more comprehensive view of the genomic regulation landscape, potentially improving the predictive power of Methven for non-coding variants.

In summary, Methven shows its potential in the functional interpretation of non-coding variants on DNA methylation, offering a more nuanced and dynamic approach to understanding their effects on DNA methylation and, by extension, on gene regulation in various diseases. As GWAS continue to expand our catalog of genetic variants, tools like Methven are essential for translating these findings into actionable biological insights, advancing the fields of genomics and personalized medicine.

## Methods

### Datasets

#### The meQTL EPIC Database

The meQTL EPIC dataset[12] was download from the meQTL EPIC Database website (https://epicmeqtl.kcl.ac.uk/), which reported the results of a meQTL analysis at 724,499 CpGs profiles in 2,358 blood samples from three UK cohorts. In this study, we obtained meQTL data from CD4+ T cells in the EPIC meQTL Database, which we used as the intra-dataset. Additionally, we utilized monocyte meQTL data from the same database as one of the external validation datasets.

#### EpiMap Repository

The corresponding ATAC-seq data for matching CD4+ T cell meQTL and monocyte meQTL were downloaded from the EpiMap Repository[13] (https://compbio.mit.edu/epimap/), which includes aggregated and uniformly re-processed functional genomics data from 3,030 references across sources such as ENCODE and Roadmap.

#### GWAS summary of RA risk SNPs

The summary results of RA risk SNPs in 101 risk loci were derived from a three-stage trans-ethnic meta-analysis, involving a genome-wide association study (GWAS) of over 100,000 subjects of European and Asian ancestries[22].

#### RA-stimulated and unstimulated ATAC-seq of CD4+ T cells

In the disease-SNP analysis, we downloaded the ATAC-seq data generated from CD4+ T cells at different time points (0 min, and 24 h) after stimulation with anti-CD3/anti-CD28 which contains information about chromatin dynamics associated with rheumatoid arthritis (RA)[23].

### Data pre-processing

#### Balanced sampling

For each SNP, CpG sites within a 100kbp range upstream and downstream that exhibit methylation level changes were identified. SNP-CpG pairs with distances less than 10kbp formed the small dataset, while those with distances between 10kbp and 100kbp comprised the large dataset. To prevent model bias, we balanced the dataset by down-sampling the more prevalent classification category according to chromosome distribution, ensuring that the number of positive and negative samples was approximately equal (Table S1). This balanced dataset was then used as the intra-dataset for both the classification and regression tasks.

#### Fetch initial input feature

For each SNP-CpG pair, DNA sequences and corresponding ATAC-seq data were extracted with the CpG site at the center, spanning a radius of 10kbp or 100kbp, ensuring the SNP falls within this range. The sequences, both pre- and post-mutation, were subjected to positional-wise cutting centered on the CpG site, and the resulting fragments were input into the DNA language model to generate DNA embeddings.

#### Embedding generation

Due to the maximum input length constraint of DNABert2, the DNA sequences must be segmented. By centering each cut on the CpG site—an essential point of interest—this segmentation ensures that each fragment remains within the permissible input length of the DNA language model while maximizing the retention of critical information integrity.

The positional-wise cutting algorithm can be described as follows: given a DNA sequence/ATAC-seq centered on a CpG site (with a total length of 10,001bp for small pairs and 100,001bp for large pairs), the first and last cuts are 250bp in length, while a fragment centered on the CpG site, with a length of 501bp, forms the central cut. The remaining DNA sequence between the terminal cuts and the central cut is uniformly segmented into cuts of 500bp each. Specifically, the DNA sequence between the first cut and the central cut is divided into n cuts of 500bp, with the same applied to the sequence between the central cut and the last cut. For the small model, n equals 19, and for the large model, n equals 199 (Fig. 1b).

Finally, the DNA embeddings from each cut and the ATAC-seq data underwent average pooling and were concatenated.

### Benchmarking

#### Definition of classification and regression task

In this study, we designed two independent tasks: a classification task to predict the direction of the SNP’s impact on CpG methylation levels (up-regulation/down-regulation) and a regression task to predict the magnitude of this impact (slope). In the classification task, samples with a slope greater than 0 were defined as positive samples (up-regulation), while those with a slope less than 0 were defined as negative samples (down-regulation). This layered prediction approach allows Methven to focus on each predictive goal independently, minimizing task interference and achieving higher predictive accuracy, particularly in determining the direction of the methylation impact.

#### Evaluation metrics

For the classification task, we employed Accuracy (ACC), Precision, Recall, F1-score, and Area Under the Curve (AUC) to assess model performance. For the regression task, we utilized Root Mean Square Error (RMSE), R-Squared and Pearson Correlation Coefficient (PCC) as key metrics.

#### Dataset split

In this study, the intra-dataset was divided using two distinct strategies. The first strategy was designed to facilitate model performance comparisons and was structured to support ten-fold cross-validation. The second strategy was applied for finalizing and releasing the model, involving a random split of the data into training, validation, and testing sets in an 8:1:1 ratio.

### Training details of Methven

Methven was built and trained using the TensorFlow 2.7 library on an NVIDIA 3090 GPU (Table S2). For the classification task, the model was trained using binary cross-entropy as the loss function, while mean squared error (MSE) was employed for the regression task. All models were trained with a learning rate of 0.001, utilizing the Adam optimizer, and an early stopping strategy with a patience of 10 epochs was applied.

### Comparison methods

#### CpGenie

To compare Methven with the current state-of-the-art tool for predicting the impact of non-coding mutations on methylation, we implemented CpGenie following the official guidelines (https://github.com/gifford-lab/CpGenie). Since the official version of CpGenie only supports predictions for CpG sites within a 500bp range upstream and downstream of SNPs, we extended the CpGenie model by expanding the input dimensions while maintaining the identical convolutional layers and hyperparameters (such as the number of filters, kernel size, and stride) as in the original version. The extended CpGenie was trained using the ten-fold cross-validation dataset, and an early stopping strategy with a patience of 10 epochs was applied (same as Methven). We extracted the output from the layer preceding the final output layer for each fold, representing the highest-level features learned by the CpGenie model. The extracted representations from each fold were then fed into a decision tree implemented using the scikit-learn library, with default parameters, to evaluate the performance. For a fair comparison of representation learning capabilities, we applied the same process to Methven.

#### Enformer

Enformer is capable of providing functional annotations for DNA sequences within approximately a 10kbp range upstream and downstream of SNPs. We obtained these annotations using the official running interface (https://github.com/google-deepmind/deepmind-research/tree/master/enformer) and applied them to the ten-fold cross-validation dataset. For each fold, the extracted functional annotations were also fed into a decision tree implemented with default parameters using the scikit-learn library. By comparing Methven with Enformer, we aimed to evaluate how the representations learned by Methven perform relative to functional annotations in predicting the impact of non-coding mutations on methylation.

### External validation on monocyte dataset

The processing of monocyte meQTL data followed the same procedure as that used for the intra-dataset. During end-to-end training, the Methven model architecture employed was identical to that used for training on CD4+ T cells, with all model weights initialized randomly. In the fine-tuning process, however, the initial weights of the monocyte model were derived from the Methven model trained on CD4+ T cells. Regardless of whether end-to-end training or fine-tuning was applied, the training details for the monocyte model were consistent with those used for the CD4+ T cell model.

### Joint analysis with disease-associated SNPs

#### Definition of affect and unaffected CpG sites

We used the Methven regression model to predict the impact of both case and control SNPs along with their corresponding ATAC-seq data. Subsequently, based on the differences in the predicted slopes (i.e., the predicted slope for the case minus the predicted slope for the control), all annotated CpG sites were categorized into four groups: Up-enhanced (where methylation impact is positively enhanced by disease occurrence), Down-enhanced (where methylation impact is negatively enhanced), Reduced (where methylation impact decreases), and Unaffected (where methylation impact remains unchanged by disease occurrence). The specific calculation can be defined by the following formula:

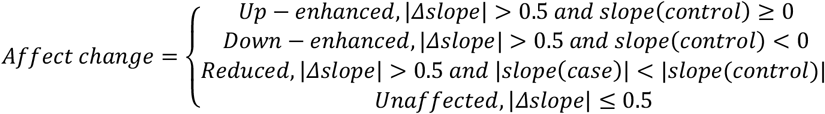

#### ATAC-seq data pre-processing

ATAC-seq data from unstimulated (0h) and 24-hour stimulated samples were used as control and case ATAC-seq, respectively. The downloaded ATAC-seq data were converted to bigwig files using BEDTools[27] and BedGraphToBigWig[28], and subsequently aligned to the hg19 genome using CrossMap[29]. These data were matched with DNA sequences corresponding to RA risk SNPs and then input into Methven for prediction.

#### GO enrichment and analysis

We calculate the distance from each CpG site to its nearest gene and annotate the distance to the gene’s transcription start site (TSS). CpG sites are then filtered based on an absolute difference in predicted slopes of 0.5 or greater, indicating significant changes in methylation levels. We further focus on CpG sites located within 2kb of the TSS, as these sites are in the proximal promoter region and may directly influence transcription initiation and gene expression regulation. The filtered CpG sites are then used to identify neighboring genes for pathway enrichment analysis using Metascape[30].

## Data availability

The GRCh37/hg19 genome and functional annotation was obtained from UCSC Genome Browser (https://genome.ucsc.edu/). The CpG annotation of Infinium MethylationEPIC v1.0 B5 manifest file was downloaded from https://support.illumina.com/downloads/infinium-methylationepic-v1-0-product-files.html. The CD4+ T cell ATAC-seq data was obtained from EpiMap Repository with BSS number ‘BSS01347’, and the monocyte ATAC-seq was downloaded using ‘BSS01279’. The 24h RA-stimulated and unstimulated ATAC-seq of CD4+ T cells are accessible through GEO Series accession number GSE138767. The GWAS summary of RA risk SNPs used in this study was obtained at http://plaza.umin.ac.jp/~yokada/datasource/software.htm with ‘Summary results of RA risk SNPs in 101 risk loci’. The data used for training, evaluating, and visualizing Methven can be downloaded on GitHub (https://github.com/Liuzhe30/Methven)

## Code availability

The source code of the pre-processing, Methven modelling and validation processes are freely available on GitHub (https://github.com/Liuzhe30/Methven) with detailed instructions.

