## Supplementary Materials for "Predicting the effect of non-coding mutations on single-cell DNA methylation using deep learning"

**Supplementary Information**

### Tables

Table S1. The data distribution of the Methven intra-data.

| **Model size** | **type** | **No. of Up-regulation pairs** | **No. of Down-regulation pairs** | **No. of pairs (all)** |
| --- | --- | --- | --- | --- |
| small | train | 7,959 | 7,940 | 15,899 |
| small | valid | 1,016 | 971 | 1,987 |
| small | test | 962 | 1,026 | 1,988 |
| large | train | 12,151 | 12,101 | 24,252 |
| large | valid | 1,503 | 1,529 | 3,032 |
| large | test | 1,534 | 1,298 | 3,032 |

Table S2. Training details of Methven.

| **Model size** | **BiGRU layers** | **BiGRU units** | **Training GPU** | **Batch size** | **No. of model hypermeters** |
| --- | --- | --- | --- | --- | --- |
| small | 2 | 64 | 1*Nvidia 3090 | 1,024 | 722,786 |
| large | 2 | 64 | 1* Nvidia 3090 | 512 | 814,946 |

Table S3. Ablation study of Methven-small model in classification.

| **Model** | **ACC** | **Precision** | **Recall** | **F1-score** | **AUC** | **No. of model hypermeters** |
| --- | --- | --- | --- | --- | --- | --- |
| Without ATAC-seq | 0.8828 | 0.8777 | 0.8805 | 0.8791 | 0.9389 | 722,402 |
| Without DNABert2 | 0.5000 | 0.4891 | 0.7672 | 0.5973 | 0.5035 | 132,962 |
| Replace DNABert2 with One-Hot | 0.9095 | 0.8924 | 0.8920 | 0.9070 | 0.9548 | 5,245,794 |
| Replace BiGRUs with FC layers | 0.8778 | 0.8772 | 0.8690 | 0.8731 | 0.9424 | 126,882 |
| Methven-small | 0.9120 | 0.9174 | 0.9063 | 0.9116 | 0.9654 | 722,786 |

Table S4. Ablation study of Methven-large model in classification.

| **Model** | **ACC** | **Precision** | **Recall** | **F1-score** | **AUC** | **No. of model hypermeters** |
| --- | --- | --- | --- | --- | --- | --- |
| Without ATAC-seq | 0.8087 | 0.8210 | 0.7953 | 0.8079 | 0.8872 | 814,562 |
| Without DNABert2 | 0.5129 | 0.5098 | 0.7761 | 0.6474 | 0.5700 | 225,122 |
| Replace DNABert2 with One-Hot | 0.8174 | 0.8269 | 0.8277 | 0.8248 | 0.9065 | 51,325,794 |
| Replace BiGRUs with FC layers | 0.8183 | 0.8377 | 0.7705 | 0.8027 | 0.8935 | 219,042 |
| Methven-large | 0.8480 | 0.8463 | 0.8530 | 0.8490 | 0.9271 | 814,946 |

Table S5. Comparison of existing methods in classification.

| **Model size** | **Model** | **ACC** | **Precision** | **Recall** | **F1-score** | **AUC** |
| --- | --- | --- | --- | --- | --- | --- |
| small | Methven | 0.9077±0.0091 | 0.9090±0.0090 | 0.9063±0.0145 | 0.9076±0.0093 | 0.9077±0.0091 |
|  | CpGenie* | 0.7818±0.1902 | 0.7399±0.2975 | 0.8261±0.2796 | 0.7656±0.2730 | 0.8067±0.2063 |
|  | Enformer | 0.8127±0.0122 | 0.8141±0.0173 | 0.8116±0.0144 | 0.8125±0.0129 | 0.8127±0.0122 |
| large | Methven | 0.8416±0.0089 | 0.8406±0.0122 | 0.8441±0.0058 | 0.8424±0.0073 | 0.8417±0.0089 |
|  | CpGenie* | 0.7057±0.1706 | 0.6564±0.2670 | 0.8076±0.2780 | 0.7071±0.2486 | 0.7388±0.1975 |
|  | Enformer | 0.7554±0.0088 | 0.7544±0.0149 | 0.7593±0.0135 | 0.7568±0.0100 | 0.7555±0.0088 |

* Predictions beyond the scope of CpGenie use a retrained CpGenie model.

Table S6. External analysis on monocyte meQTLs.

| **Model size** | **Model** | **ACC** | **Precision** | **Recall** | **F1-score** | **AUC** |
| --- | --- | --- | --- | --- | --- | --- |
| small | end-to-end | 0.845 | 0.802 | 0.890 | 0.843 | 0.898 |
|  | fine-tune | 0.888 | 0.868 | 0.898 | 0.883 | 0.939 |
| large | end-to-end | 0.704 | 0.665 | 0.825 | 0.736 | 0.770 |
|  | fine-tune | 0.763 | 0.742 | 0.809 | 0.774 | 0.823 |

Table S7. Prediction results of Methven-small on rs968567 (chr11: 61595564) in disease-SNP analysis.

| **CpG site** | **CpG position** | **CpG-SNP distance** | **Slope(0h)** | **Slope(24h)** |
| --- | --- | --- | --- | --- |
| cg07709195 | 61586015 | 9,549 | 3.30768 | 3.107404 |
| cg08281583 | 61595223 | 341 | -0.97285 | 0.414493 |
| cg02563962 | 61595550 | 14 | 0.895135 | -0.17976 |
| cg20896974 | 61595983 | 419 | 1.704718 | 0.675927 |
| cg15454066 | 61595377 | 187 | -0.55628 | -0.67076 |
| cg21409469 | 61594769 | 795 | -1.50895 | 0.143971 |
| cg07005513 | 61595956 | 392 | 1.896686 | 1.641777 |
| cg19481605 | 61596812 | 1,248 | -2.00758 | -0.96298 |
| cg06781209 | 61594997 | 567 | -1.18766 | 0.085661 |
| cg14911132 | 61596755 | 1,191 | 2.173507 | 3.008933 |
| cg05698098 | 61595494 | 70 | -1.24906 | -1.56824 |
| cg27386326 | 61587980 | 7,584 | -3.43851 | -2.88585 |
| cg25324164 | 61598330 | 2,766 | 0.943897 | -0.18036 |
| cg13299762 | 61594708 | 856 | -2.75586 | -1.79238 |
| cg08380661 | 61592206 | 3,358 | 2.004973 | 1.569955 |
| cg07999042 | 61598930 | 3,366 | -2.5631 | -2.4589 |
| cg23760165 | 61595485 | 79 | -0.09937 | -0.28168 |
| cg14562930 | 61595050 | 514 | -1.89603 | -1.14012 |
| cg05816884 | 61595492 | 72 | -0.91971 | -0.94432 |
| cg19610905 | 61596333 | 769 | 0.973409 | -0.29367 |
| cg08093537 | 61595225 | 339 | -0.29462 | 0.866731 |
| cg25303599 | 61595807 | 243 | 1.69671 | 1.639525 |
| cg00614641 | 61596405 | 841 | -2.71884 | -3.82376 |
| cg10515671 | 61585899 | 9,665 | 1.785255 | 1.755254 |
| cg09610223 | 61595465 | 99 | -0.13369 | -0.34711 |
| cg22796604 | 61602232 | 6,668 | -3.07965 | -3.84196 |
| cg00603274 | 61596626 | 1,062 | 0.998823 | 1.712455 |
| cg21709803 | 61594965 | 599 | -2.26192 | -0.98825 |
| cg11250194 | 61601937 | 6,373 | -2.95164 | -2.51638 |
| cg10868875 | 61596307 | 743 | -0.01807 | -1.01722 |
| cg21029357 | 61601062 | 5,498 | -1.52554 | -1.73847 |
| cg01400685 | 61598025 | 2,461 | 1.165036 | -0.20022 |

Table S8. The data distribution of the external monocyte dataset.

| **Model size** | **type** | **No. of Up-regulation pairs** | **No. of Down-regulation pairs** | **No. of pairs (all)** |
| --- | --- | --- | --- | --- |
| small | train | 2,014 | 2,000 | 4,014 |
| small | valid | 259 | 243 | 502 |
| small | test | 236 | 266 | 502 |
| large | train | 3,371 | 3,377 | 6,748 |
| large | valid | 420 | 423 | 843 |
| large | test | 423 | 421 | 844 |

### Figures

Figure S1. The data distribution of the Methven intra-data in pre-processing progress.

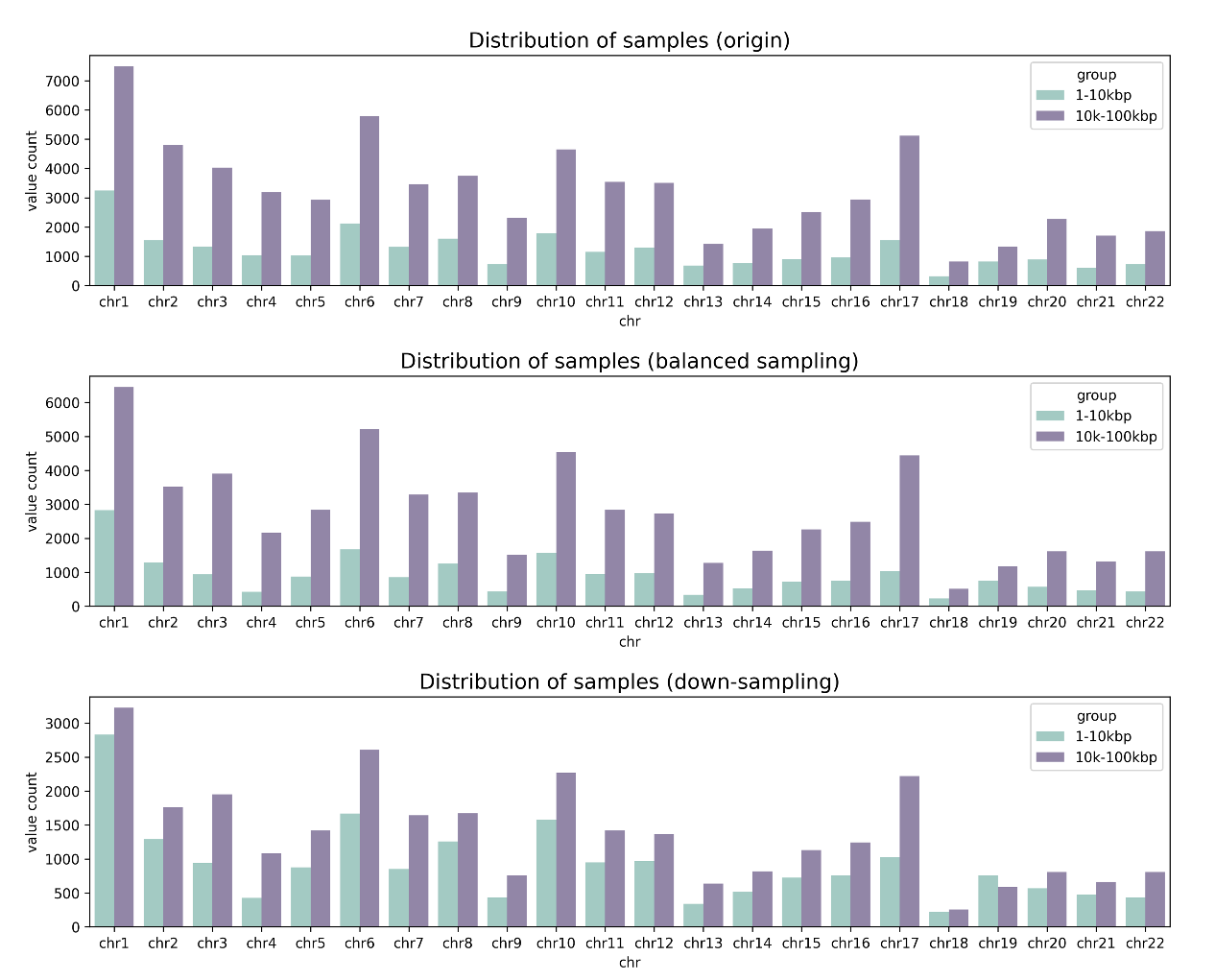

Figure S2. The data distribution of the external monocyte dataset in pre-processing progress.

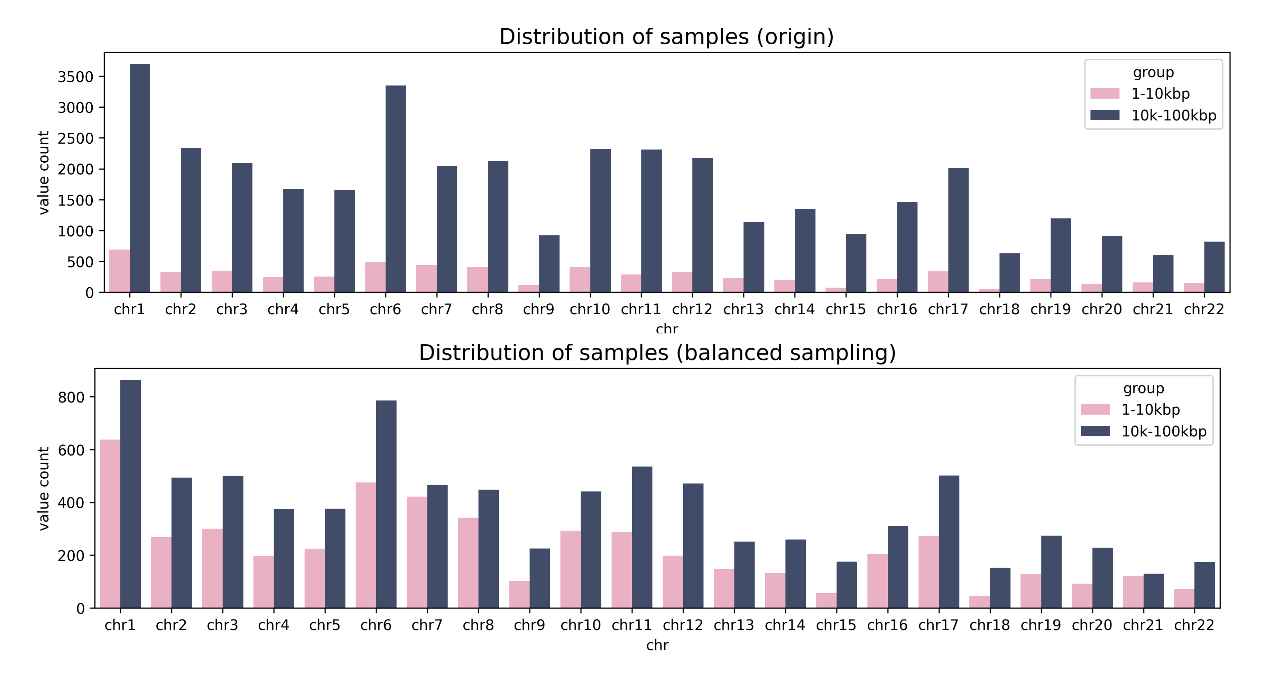

Figure S3. The performance of existing methods in model comparison.

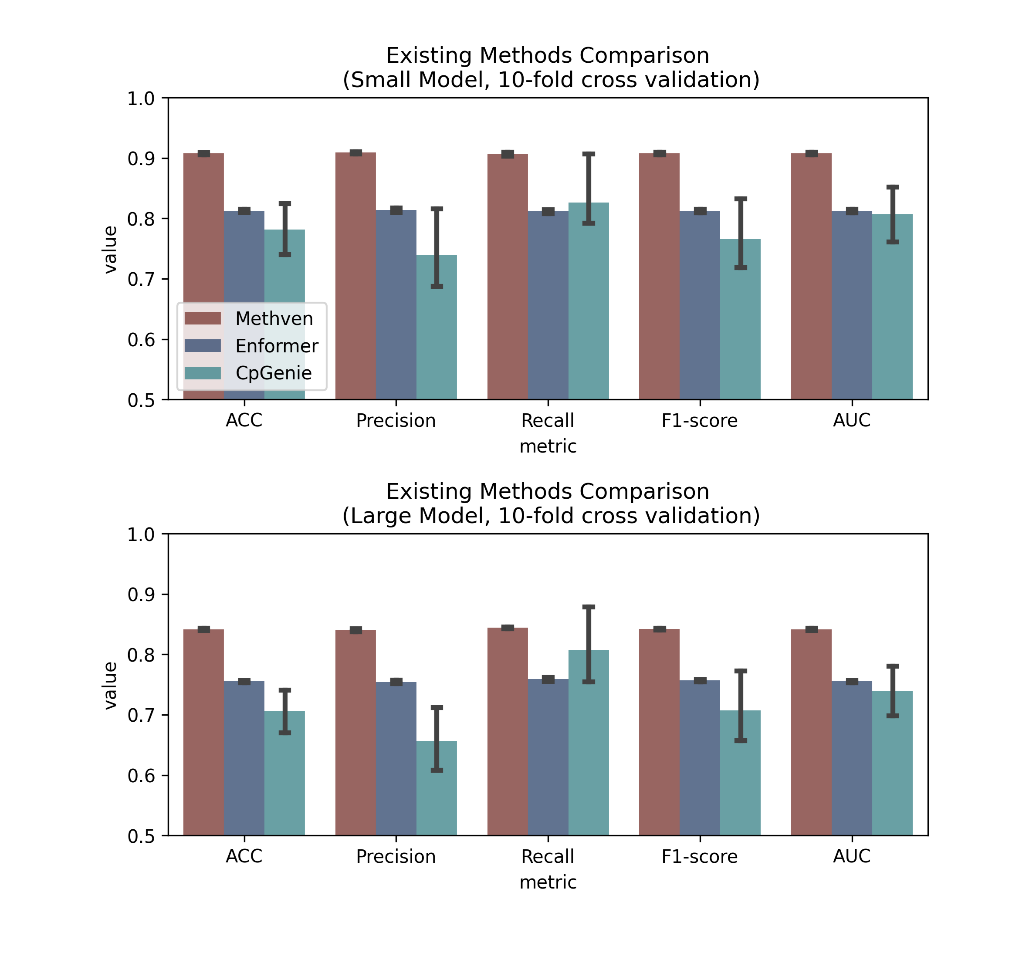
